# Rescue by gene swamping as a gene drive deployment strategy

**DOI:** 10.1101/2022.03.08.483503

**Authors:** Keith D. Harris, Gili Greenbaum

**Affiliations:** Department of Ecology, Evolution and Behavior, The Hebrew University of Jerusalem, Jerusalem, Israel

## Abstract

Gene drives are genetic constructs that can spread deleterious alleles with potential application to population suppression of harmful species. Given that a gene drive can potentially spill over to other populations or even other species, control measures and fail-safes strategies must be considered. Gene drives are designed to generate a rapid demographic decline, while at the same time generating a dynamic change in the population’s genetics. Since these evolutionary and demographic processes are linked and are expected to occur at a similar time-scale during gene drive spread, feedback between these processes may significantly affect the outcome of deployment. To study this feedback and to understand how it affects gene drive spillovers, we developed a gene drive model that combines evolutionary and demographic dynamics in a two-population setting. The model demonstrates how feedback between evolutionary and demographic dynamics can generate additional outcomes to those generated by the evolutionary dynamics alone. We identify an outcome of particular interest, where the short-term suppression of the target population is followed by gene swamping and loss of the gene drive. This outcome could be useful for designing gene drive deployments that temporarily suppress the population, but ultimately do not remain in the population. Using our model, we demonstrate the robustness of this outcome to spillover and to the evolution of resistance, and suggest that it could be used as a fail-safe strategy for gene drive deployment.

## 1 Introduction

Gene drives are genetic constructs that allow the spread of deleterious alleles by violating Mendelian inheritance patterns [1]. With CRISPR/Cas9-based technology, gene drives can potentially be used to suppress or eliminate wild populations [2, 3], and have the potential for revolutionizing biocontrol [4]. However, there are substantial concerns that need to be addressed before deployment in the wild can be considered, including the spillover of the gene drive to other populations or species [5, 6], and the evolution of resistance to the gene drive [7]. Addressing these concerns is complicated by the potential interaction between the evolutionary dynamics of the spread of the gene drive in the population and its demographic impacts. In potential gene drive deployments, the evolutionary and demographic effects are expected to occur at similar time scales, and so feedback between these processes may substantially shape the outcome [8]. Therefore, in order to inform gene drive design and to develop novel strategies for mitigating the risks of gene drive deployments, investigating the evolutionary-demographic dynamics of gene drives is crucial.

Mathematical and computational modeling has been instrumental in guiding gene drive research, allowing the study of key aspects in the behavior of gene drives prior to their deployment [9, 10]. Population genetic models investigating the evolutionary dynamics of non-Mendelian inheritance were developed decades before the recent development of CRISPR-based drives [11, 12], and were subsequently adjusted to more specifically model CRISPR-based gene drives [13, 14]. Some models focus on whether gene drives could effectively spread in a single-population [12–14], while others focus on spread of the gene drive between discrete populations [6, 15] or across a continuous landscape [8, 16, 17]. These models showed that, in cases in which the gene drive spreads in a population or region regardless of its initial frequency, it will spill over to all other connected populations or regions. Therefore, only with threshold-dependent gene drives (i.e., cases where the gene drive spreads only when its frequency is above a critical threshold) are spillovers potentially avoidable. Modeling efforts have also been directed to studying the evolution or resistance to the gene drive constructs [18], showing that in some cases resistance alleles are expected to spread rapidly in a population once they appear [19], leading to the loss of the gene drive allele, and increasing the chance of future deployments failing.

The consequence of feedback from the demographic changes caused by the gene drive on the evolutionary dynamics has also been studied in continuous-space models, and has been shown to affect the outcome of gene drive deployment [8, 20]. In these models, the decrease in local density caused by the negative demographic effect of the gene drive allele can prevent the gene drive from spreading, or lead to long-term persistence of the gene drive with fluctuations of both the gene drive allele frequency and local population density. While these models demonstrate that demography can crucially affect the success of deployment in a single population, it is unclear how demographic feedback would affect the spillover of the gene drive from the target population to other, non-target, populations. In particular, demographic changes could lead to asymmetrical gene flow between the target population and neighboring populations, and thus substantially alter the risk of spillover [6].

In order to study how the interaction between the evolutionary and demographic processes involved in gene drive spread influences deployment outcomes, we developed a model that tracks interconnected evolutionary and demographic dynamics. For tractability, the model is set in a two-population system, which allows for spillovers from the target population to the non-target population, as well as for potential gene swamping of the target population by wild-type alleles from the non-target population. Our investigation is specifically focused on identifying emerging outcomes which could be used to develop strategies that can mitigate the risks of gene drive spillovers, and reduce the probability of resistance evolution.

## 2 Model

To study gene drive dynamics and spillovers in the presence of feedback between the evolutionary dynamics and the demographic impact of the gene drive, we developed and investigated two-population mathematical models tracking the gene drive allele frequency and the population sizes. Our focus was to investigate conditions under which the gene drive is potentially able to spread to different frequencies in the two populations (“differential targeting” gene drives [6]), and therefore we limited our study to suppression gene drives [4], in which the fitness cost of the gene drive allele can generate threshold-dependence [14]. Modification gene drives, which do not substantially affect fitness, cannot be threshold-dependent, and are expected to result in spillover to populations connected by gene flow whenever the gene drive spreads in a single population; we do not study such gene drives here.

Our model consists of two dynamics that are related to the spread of gene drives: (i) an evolutionary dynamic, tracking the change in the gene drive allele frequency *q*, and (ii) a demographic dynamic, tracking the impact of the gene drive on populations sizes *N*. In order to understand the impact of gene flow during gene drive spread, we model two populations, identical in terms of their carrying capacity, which are connected by migration. We formulate two sets of recursive equations, one set for tracking the gene drive allele frequency in the two populations *q*_*i*_, and one set for tracking the two population sizes, *N*_*i*_(*i* = 1, 2). For simplicity and coherence, we define population 1 as the ‘target population’ that we aim to suppress and where the gene drive is deployed, and population 2 as the ‘non-target population’. The model is implemented and available on the modelRxiv platform [21], where all results can be regenerated and the model can be re-parameterized (https://modelrxiv.org/model/yoKkSv).

### 2.1 Migration between populations

We consider the case where migration occurs prior to selection, and we model changes in population sizes in two steps: from pre-migration to post-migration, and then from post-migration to post-selection. We consider here only the migration-before-selection case because the results are typically similar in the selection-before-migration case [6]. For simplicity, we only consider the case where migration is inherently symmetric, with a proportion *m* of each population migrating per generation (i.e., symmetric in terms of proportion of migration, but may be asymmetric in terms of number of individuals migrating when the population sizes are unequal). We formulate the change in the population size of population *i* from the pre-migration sizes, denoted as *N*_*i*_, to the post-migration sizes, denoted as *Ñ*_*i*_:

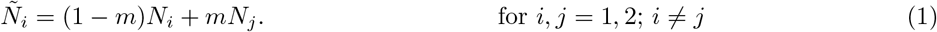

To model the impact of migration on the genetic composition of population *i*, we formulate the change from the pre-migration gene drive allele frequency in population *i*, denoted as *q*_*i*_, to the post-migration frequency, denoted as 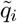:

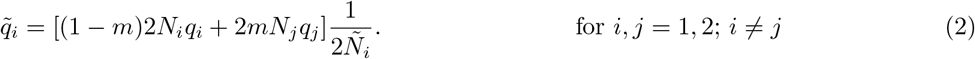

In Eq. 2, the number of gene drive alleles in the population following migration is normalized by the population size to compute the post-migration allele frequencies.

### 2.2 Evolutionary dynamics

To model evolutionary dynamics, we use a previously studied model of gene drive dynamics in two populations [6], which is based on a model of gene drive spread in a single population [13, 14]. This model considers the fitness cost of the gene drive homozygote relative to the wild type, *s*, the dominance of the gene drive allele, *h*, and the conversion rate of heterozygotes to gene drive homozygotes by the CRISPR copying mechanism, *c*. In our model we consider gametic conversion of the wild type allele (we present results for a zygotic conversion model in Supplementary Information 1). The change in the gene drive allele frequency in population *i* from the post-migration frequencies 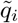 (Eq. 2) to the post-selection frequencies 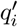 can be described by the recursion equation [6]

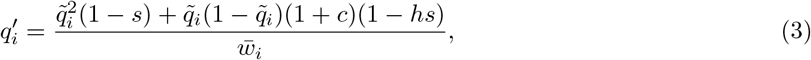

where the average fitness 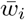 is defined as 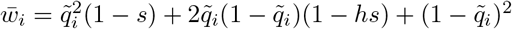

### 2.3 Demographic dynamics

To model demographic dynamics, we use the classic logistic equation to describe changes in populations sizes following migration (*Ñ*_*i*_ in Eq. 1):

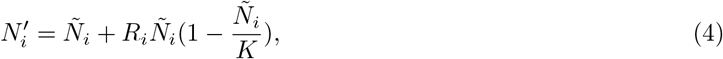

where *K* is the carrying capacity (assumed identical for both populations) and *R*_*i*_ is the growth rate of population *i*. We also consider the populations identical in terms of their intrinsic growth rates. However, as the growth rate is affected by the spread of the gene drive, we treat the actual growth rate of the populations as a dynamic parameter, *R*_*i*_, which depends on the gene drive allele frequency. To describe this dependency, we denote the intrinsic growth rate in the absence of the gene drive as *r*_0_, and the growth rate of a population fixed for the gene drive allele as *r*_1_. For heterozygotes, we assume a growth rate proportional to the dominance parameter *h*, i.e., *r*_*h*_ = (1 *− h*)*r*_0_ + *hr*_1_. The population growth rate *R*_*i*_ at a given point in time is, therefore, computed by weighing the different growth rates with the Hardy-Weinberg genotype frequencies of the gene drive allele:

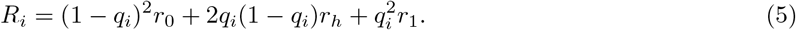

In this equation the contribution of the heterozygote genotype frequencies is different for converted and non-converted genotypes, as in Eq. 2. Eq. 5 links the demographic dynamics described in Eqs. 1 and 4 with the evolutionary dynamics of Eqs. 2 and 3, which together comprise the dynamical system we study.

### 2.4 Demographic effect of the gene drive

The parameter *r*_1_ describes the demographic impact of the gene drive, and is related to the relative fitness cost *s* of the gene drive allele. The relationship between *r*_1_ and *s* is of particular interest, because it defines the population-level impact of the gene drive allele, and therefore could vary depending on the demographic effect of the gene drive phenotype on the population. If the gene drive has no demographic effect (i.e., gene drive carriers have decreased intra-species competitive abilities, but do not suffer from absolute fitness costs), *r*_1_ should be independent of *s* and equal to the intrinsic growth rate *r*_0_. If the demographic effect is correlated to the effect of the gene drive on relative fitness, *s* and *r*_1_ should be tightly linked. To model this relationship, we introduce a parameter *d* that allows the manipulation of the level of the demographic effect of the gene drive; this parameter determines the degree to which growth rate *r*_1_ is impacted in relation to the relative fitness cost *s*:

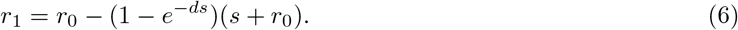

Under this formulation, for *d* = 0, the gene drive has no effect on growth rate (i.e. soft selection; *r*_1_ = *r*_0_), whereas as *d* → ∞, we have *r*_1_ → −*s*, meaning that the (negative) growth rate is fully correlated with the relative fitness cost. Intermediate values of *d* represent partial correlations between within-population competition and population-level demographic effects. The formulation in Eqs. 6 was chosen to allow the manipulation of the demographic effect of the gene drive and generate a continuum between the purely evolutionary dynamics and the demographic-evolutionary dynamics, and does not represent any particular relationship between genotypes and fitness. An alternative simplified formulation of *r*_1_ does not substantially alter the outcome of the model (see section ‘Effect of *r*_1_ formulation on deployment outcomes’ in Supplemental Information).

## 3 Results

### 3.1 The effect of demographic-evolutionary feedback on outcomes

To understand how demography influences the outcome of gene drive deployment, it is insightful to compare the outcomes of our model to that of a previous model taking into account only evolutionary dynamics [6], which is equivalent to *d* = 0 in our model. In the purely evolutionary model, three types of qualitatively different outcomes are expected following deployment in the target population: (i) the gene drive deployment fails and the gene drive allele is lost from both populations (‘*failure*’); (ii) the gene drive spills over from the target population to the non-target population and is driven to fixation in both (‘*spillover*’); or (iii) the gene drive remains in a stable state (high frequency in the target population and low frequency in the nontarget population) for low migration rates (‘*differential targeting*’), but spillover or failure occurs for higher migration rates. The outcome of deployment is determined by the gene drive configuration parameters (*s, c*, and *h*), and for gene drive designs that are threshold-dependent (in a single population model), also by the migration rate, *m*. Through characterization of the dynamics under different paramaterization of our model, we observe that incorporating demographic effects of the gene drive generates additional types of outcomes (Fig. 1). We classified the types of outcomes according to the gene drive allele frequency and the demographic changes in the populations (Table S1).

**Figure 1:**
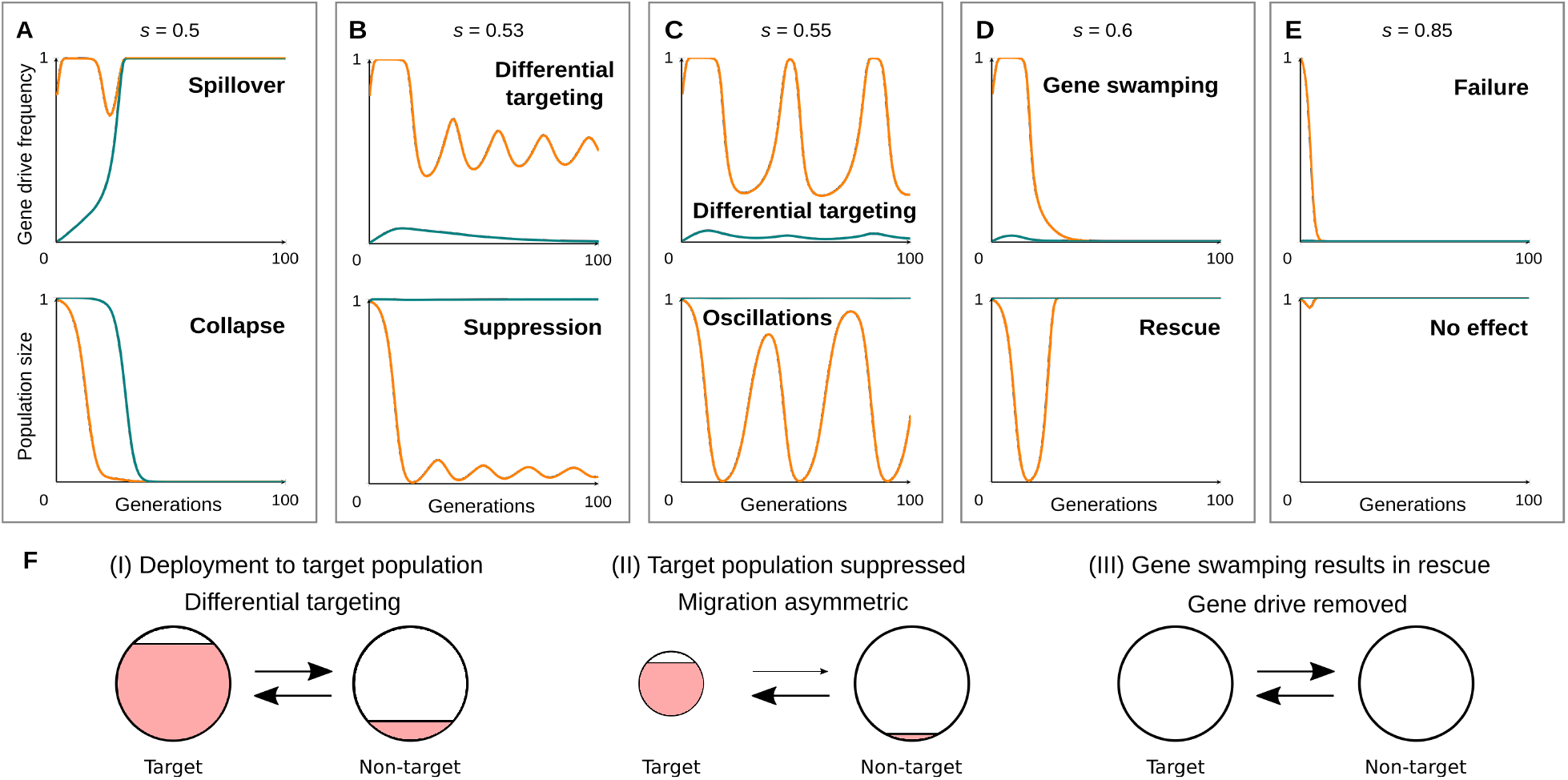
Distinct outcomes of deployment in a gene drive model incorporating feedback between evolution and demography. (A–E) Outcomes classified as indicated in Table S1. The top panels in (A–E) show the evolutionary dynamics in terms of the gene drive allele frequency, while the bottom panels show demographic dynamics in terms of the population sizes. Dynamics in the target population are shown in orange, and in the non-target population in blue. (A) Gene drive spillover followed by collapse of both populations. (B) Differential targeting with stable suppression of only the target population. (C) Oscillations in the evolutionary and demographic dynamics. (D) Suppression of the target population followed by gene swamping and global removal of the gene drive. (E) Failure of the gene drive with no demographic effect. The outcomes differ only in the relative fitness cost of the gene drive, *s*, noted above each panel, and are identical in all other parameters: *c* = 1, *h* = 1, *m* = 0.01, *q*_1_(0) = 0.8, and *d* = 10. (F) Schematic illustration of the dynamics in the gene swamping outcome in (D): (i) the gene drive is initially deployed to the target population and has a low frequency in the non-target population (gene drive allele frequency illustrated as the filled red portion); (ii) as the target population is suppressed, migration becomes increasingly asymmetric, with decreased gene flow out of and increased gene flow into the target population; (iii) gene swamping due to asymmetric migration leads to gene swamping and removal of the gene drive from both populations.

Two of the outcomes, ‘*collapse*’ (Fig. 1A) and ‘*failure*’ (Fig. 1E), are of less interest because there is no targeted demographic effect in either, and therefore they represent undesirable scenarios in terms of gene drive deployment. However, we identified three outcomes in which differential targeting of the gene drive results in distinct demographic and long-term genetic effects. The first is long-term ‘*differential targeting*’ and demographic ‘*suppression*’ of the gene drive in the target population (Fig. 1B). Here, fluctuations of the evolutionary and demographic dynamics eventually stabilize, and the system converges to a differential targeting state, as observed in purely evolutionary models that do not incorporate demography. With the other parameters remaining fixed, increasing the relative fitness cost of the gene drive *s* results in a different outcome, in which the ‘*oscillations*’ of the evolutionary and demographic dynamics do not converge, and no steady state is reached (Fig. 1C). In each cycle, a high gene drive allele frequency in the target population leads to demographic impact and a decrease in population size, as in the suppression outcome. However, consequently, gene flow from the non-target population, which now becomes proportionately larger, has an increased impact on the target population. Because this gene flow is mainly of wild type alleles (gene drive frequencies in the non-target population are low at the differential targeting state), the gene drive allele frequency in the target population is reduced. The wild-type-abundant gene flow reduces the demographic impact of the gene drive, and the target population switches from negative to positive growth. The impact of gene flow from the target population is then reduced as the target population increases in size, and the gene drive again spreads to high frequencies in the target population, ending the cycle, and a new oscillation begins.

Increasing the relative fitness cost of the gene drive further can generate an additional type of outcome (Fig. 1D), in which gene flow from the non-target population, coupled with negative selection against the gene drive allele, is sufficiently strong to completely remove the gene drive allele from the target population, resulting in ‘*gene swamping*’ [22] — complete removal of the gene drive allele — followed by full demographic recovery of the target population (Fig. 1F). In this outcome, unlike in any of the other outcomes, the gene drive allele is not expected to remain in any of the populations, while it still generates a significant demographic impact on the target population, and only on the target population. This demographic effect is not persistent as in the suppression outcome (Fig. 1B), but instead generates a short-term suppression phase that is expected to dissipate. Nevertheless, if other population control measures are introduced during the suppression phase, long-term suppression or elimination of the target population may be achieved. Therefore, to achieve control of the target species this strategy would require additional types of population controls to augment genetic suppression. However, the risk of permanent genetic impact on the population, and consequently of the evolution of resistance to the gene drive, is reduced because the expected final outcome is removal of the gene drive allele from the entire system (see section 3.5 below).

The parameter *d*, which represents the correlation between the relative fitness cost of the drive (*s*) and the growth rate of a population fixed for the gene drive allele (*r*_1_), controls the level of demographic feedback in the model. At low values of *d, r*_1_ remains positive regardless of *s* (Fig. S1A), and therefore there are no demographic changes. In this case, the model is similar to the purely evolutionary model. At higher values of *d, r*_1_ is correlated with *s* (Fig. S1A), and consequently the evolutionary dynamics of the gene drive can affect population sizes, as well as the migration rates. This results in a shift in the deployment outcome with increasing values of *d* (Fig. S1B).

### 3.2 Migration rates, gene drive configurations, and initial frequencies

To understand the relation between gene drive designs, ecological constraints and the deployment outcomes we identified, we investigated model outcomes for different gene drive configurations (values of *s, c*, and *h*) and migration rates, *m* (Fig. 2). For each migration rate value *m*, we also computed the proportion of the parameter space that each outcome occupies (for fixed *h* value, for tractability), in order to understand how difficult it would be to generate gene drive designs that can ensure a desired outcome (Fig. 2B).

**Figure 2:**
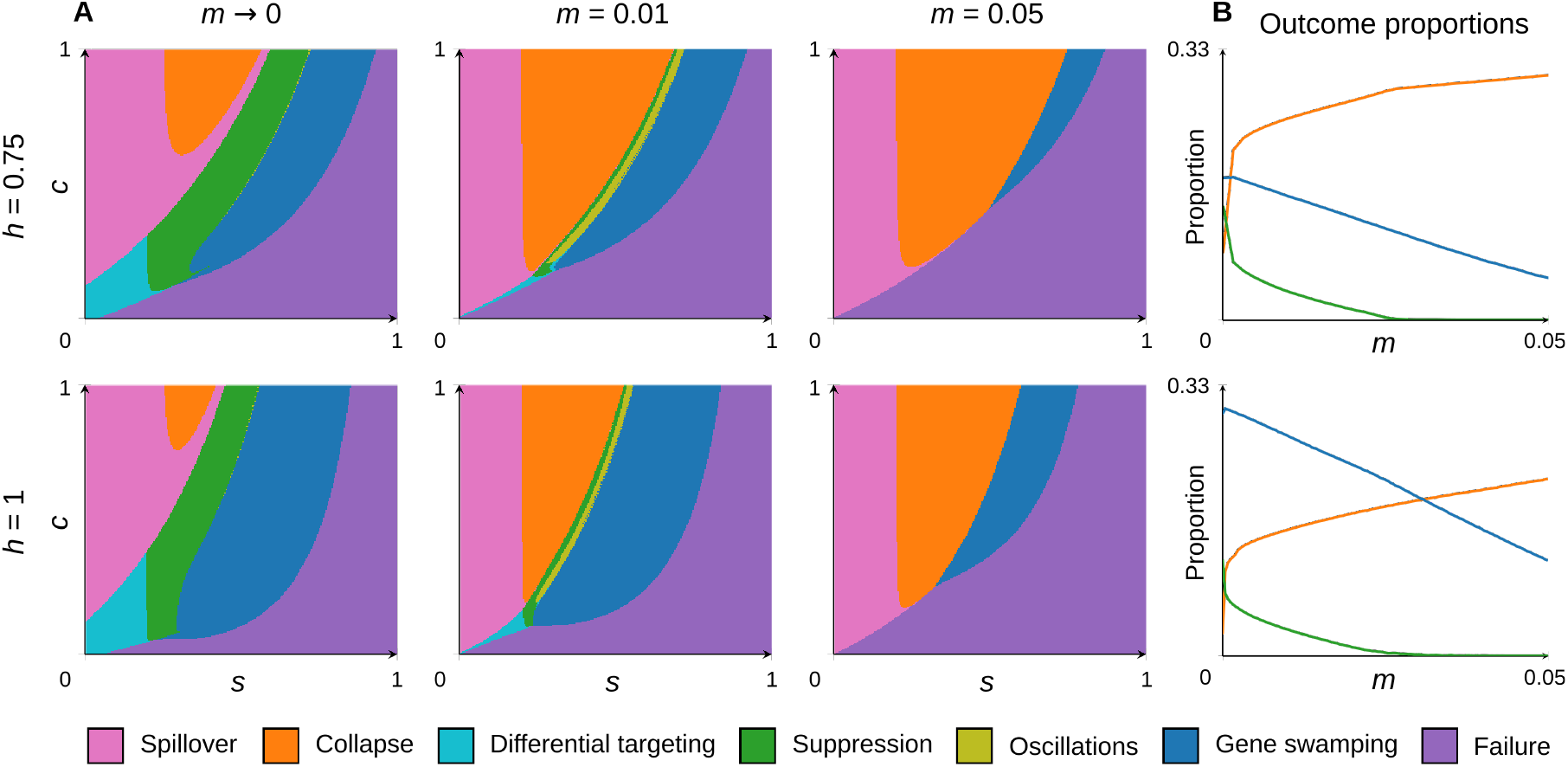
Model outcomes in relation to gene drive design and migration rates. Colors indicate the outcome as follows: ‘spillover’—spillover without collapse in pink; ‘collapse’—collapse of both populations due to spillover in orange; ‘differential targeting’—differential targeting of the gene drive without demographic effects in cyan; ‘suppression’—suppression of target population in green; ‘oscillations’—oscillations of dynamics in the target population in teal; ‘gene swamping’—gene swamping and subsequent rescue of the target population with loss of the gene drive in blue; ‘failure’—failure of the deployment (no demographic effect) and loss of the gene drive from both populations in purple. (A) Each panel shows the outcome attained in our model for a fixed *h* value, for several *m* values, and for all *s* and *c* values. (B) Relative proportions for collapse (orange), suppression (green) and gene swamping (blue) outcomes. Increasing migration rates increase the proportion of spillover outcomes, but disproportionately affect suppression outcomes. In all panels the parameters *d* = 10 and *q*_1_(0) = 0.8 were used.

For low migration rates (*m* = 1 *×* 10^*−*6^, denoted as *m →* 0; left panels in Fig. 2A) much of the parameter space results in failure or spillover of the gene drive, while, for some parameters, differential targeting, suppression or gene swamping outcomes can be achieved (particularly for higher *h* values). At higher migration rates (*m* = 0.01 and *m* = 0.05), an increased range of gene drive configurations result in spillover and collapse of both populations (orange curve in Fig. 2B). Importantly, while suppression outcomes (green in Fig. 2) quickly become almost unachievable as migration rates increase (i.e., this outcome occupies only a narrow range in the parameter space), the gene swamping outcome (blue in Fig. 2) still occupies a significant portion of the parameter range. Therefore, although the possibility of achieving gene swamping is also dependent on migration, it is achievable for higher migration rates compared to the suppression outcome (Fig. 2).

In Figures 1 and 2 we assumed a relatively high initial gene drive allele frequency in the target population (*q*_1_(0) = 0.8), meaning that four times the initial population in individuals homozygous for the gene drive allele are released. This would require an intensive release effort, comparable with some other applied population control strategies [23], but one that would be difficult in many organisms and scenarios. To assess the degree to which the outcomes we identified rely on this high initial frequency, we analyzed the proportion of three outcomes (collapse in orange, suppression in green, and gene swamping in blue) for different designs and migration rates, across a continuum of initial frequencies (Fig. S3). While the initial frequency affects the proportion of all outcomes, it affects mainly the proportion of gene drive designs allowing gene swamping (blue curves in Fig. S3). When considering only evolutionary dynamics, a higher initial frequency allows differential targeting with higher values of *s* [6]; our results indicate that these gene drive designs lead to gene swamping outcomes in a combined evolutionary-demographic model. While this result demonstrates the constraints of attaining gene swamping outcomes in terms of deployment costs, it also indicates that gene swamping is perhaps achievable even at lower initial frequencies (e.g., *q*_1_(0) *≈* 0.5; Fig. S3).

### 3.3 Gene swamping is more robust to spillover than suppression

So far we have shown that migration rates influence the proportion of outcomes, and that certain suppression outcomes are achievable only at low migration rates (Fig. 2). An important aspect of gene drive design is robustness to variability that may be present in the ecosystem, such as changing migration rates or demography, or inability to accurately estimate these parameters. In order to compare the outcomes we identified in terms of their sensitivity to an increase in migration rates, we consider gene drives with a range of fitness costs *s* with a fixed conversion rate of *c* = 1 (i.e. no heterozygotes) for different values of *m*.

When *m* is sufficiently low, it is possible to design gene drives that achieve either suppression or gene swamping (bottom part of Fig. 3A). Increasing the migration rate reduces the proportion of suppression and gene swamping outcomes (Fig. 3A), as demonstrated in Figure 2B. These outcomes, for the same parameters other than *m*, are replaced by the collapse and failure outcomes. However, the gene drive design (in this case, the value of *s*) also influences whether an increase in migration will result in spillover or failure. To illustrate this point, we consider three different designs (*s* = 0.53, 0.6, 0.7), plotted as white dashed lines in Figure 3A. The first design (*s* = 0.53) achieves suppression at low migration rates, whereas the second and third designs (*s* = 0.6 and *s* = 0.7) lead to gene swamping. Increasing the migration rate shifts the first and second designs to spillover, at different thresholds of migration. However, the third design shifts to failure when increasing the migration rate. This demonstrates that some gene drive designs that lead to gene swamping will not lead to spillover if migration rates increase. This feature, if incorporated into the design of the gene drive, could serve as a fail-safe, allowing deployment to accommodate variability in migration rate or imprecision in evaluation of migration rate, without the expected risk of spillover.

**Figure 3:**
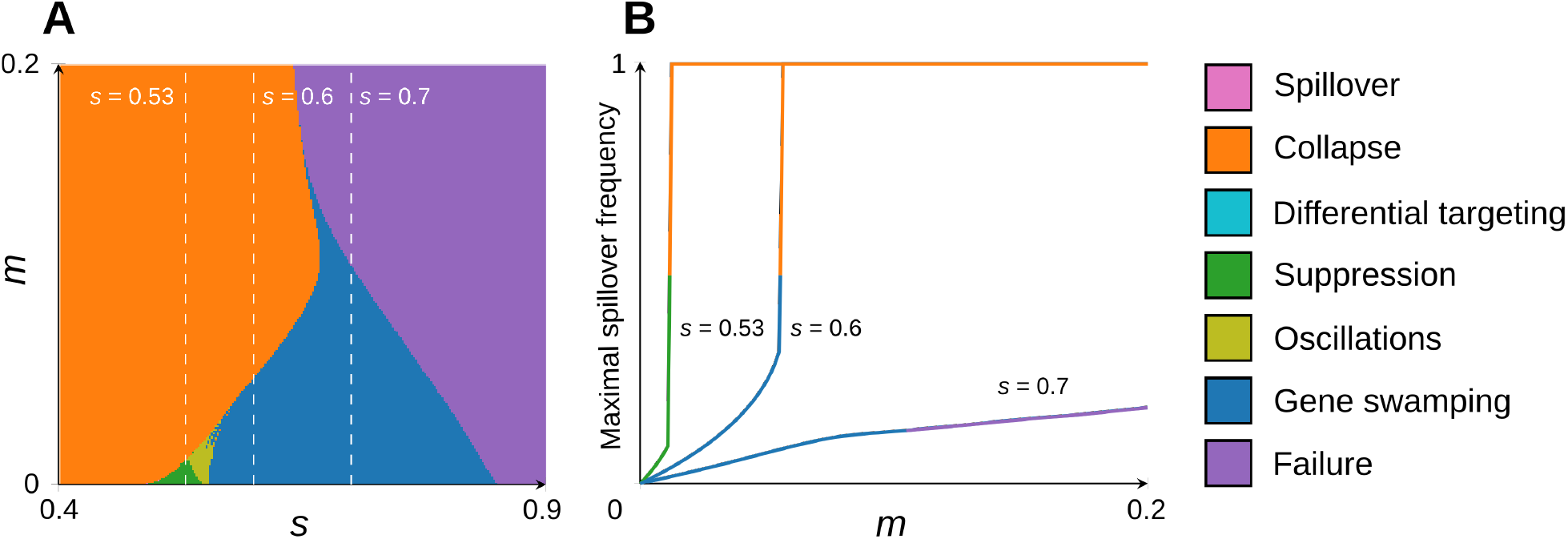
Designing gene drives to achieve ‘gene swamping’ at low migration rates and failure at high migration rates. (A) Outcomes for different fitness costs (*s*), for increasing migration rates (*m*). (B) Maximal spillover frequency (i.e., highest gene drive allele frequency in non-target population) for increasing levels of migration. Other parameters used for both panels are *c* = 1, *h* = 1, *d* = 10 and *q*_1_(0) = 0.8.

An additional aspect to gene drive deployment is the effect of the gene drive on non-target populations prior to spillover [6]; in our model, this can be measured as the gene drive frequency in the non-target population in outcomes where it is present there (suppression, oscillations, and gene swamping). This reflects not only the increased risk of spread of the gene drive in the non-target population, but also the risk of demographic effects on this population and further spread of the gene drive to other non-target populations through gene flow. For the three designs used above (*s* = 0.53, 0.6, 0.7), we also analyze the effect of deployment on the non-target population, defined as the maximal gene drive allele frequency in the non-target population during deployment, which we term the ‘maximal spillover frequency’. We plotted the maximal spillover gene drive allele frequency for these designs for increasing migration rates, with the line color indicating the deployment outcome for this migration rate (i.e., change in line color indicates a shift in outcome at a threshold level of *m* in Fig. 3B). For the first two designs that result in spillover in Figure 3A (*s* = 0.53 and *s* = 0.6), the maximal spillover frequency increases with *m* below the spillover threshold (Fig. 3B). For the third design, which does not result in spillover at higher migration rates in Figure 3A (*s* = 0.7), the maximal spillover frequency also increases with *m* but is lower than that of the other two designs.

### 3.4 Suppression characteristics in gene swamping outcomes

As opposed to long-term suppression, in the gene swamping outcome the gene drive is removed from both populations following a short-term targeted suppression phase (Fig. 1D). The feasibility of gene drives allowing short-term suppression would depend on the effectiveness of the suppression phase. In order to characterize how gene drive design and migration affect the degree of suppression in this outcome, we studied three characteristics of the suppression phase (Fig. 4A): (i) the duration (in generations) for which the target population is suppressed, (ii) the maximal level, in terms of population size or density, to which the population is suppressed, and (iii) the accumulative suppression of the target population during deployment. For gene drive designs achieving gene swamping, a higher relative fitness cost *s* results in less effective suppression, both in terms of duration and level of suppression (Fig. 4B–D). In other words, with higher values of *s* the targeted population collapses faster, shortening the suppression phase (Fig. 4B). This relationship is consistent for different migration rates, with higher migration rates reducing both the duration and level of suppression (Fig. S3).

**Figure 4:**
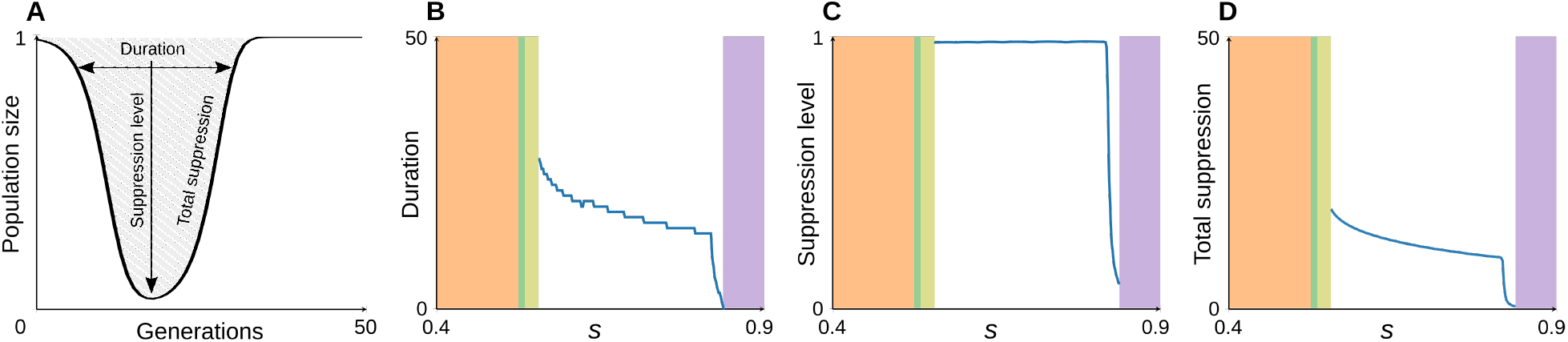
Characteristics of the ‘gene swamping’ outcome. (A) Schematic representation of the characteristics studied: ‘suppression level’ is the maximal proportion of the target population suppressed, ‘duration’ is the number of generations for which the target population size is below the threshold level *N <* 0.9*K*, and ‘total suppression’ is the accumulative suppression (area of the gray zone above the curve). (B-D) Analysis of characteristics of suppression in gene swamping outcome for different gene drive designs (different values of *s* with a fixed conversion rate *c* = 1). Blue curves indicate the value of each parameter for gene swamping outcomes; values of *s* leading to other outcomes are colored according to the color legends in Figures 2 and 3. (B) The number of generations in which the target population is suppressed (‘duration’). (C) The maximal suppression level. (D) The total suppression during gene swamping. In all panels the parameters used are *c* = 1, *h* = 1, *m* = 0.01, *d* = 10 and *q*_1_(0) = 0.8.

### 3.5 Gene swamping is robust to the evolution of resistance

One of the main challenges in gene drive deployment is to reduce the possibility of evolution of resistance to the gene drive [18, 19, 24, 25]. Resistance alleles can prevent fixation of the gene drive and even lead to its loss by competition with the gene drive allele. Resistance can arise through mutation of the wild type allele targeted by the gene drive, in the same locus, interfering with the CRISPR targeting mechanism [12, 26], or through mutation of other loci that reduce conversion efficacy [18]. The emergence and spread of resistance alleles has long-term implications, because the persistence of a resistance allele after deployment increases the chance that future deployments of the same gene drive will fail.

To evaluate the probability of resistance evolving in our model, we integrated an evolutionary model of gene drive resistance [19] into the genetic element of our model (see *Methods* section). We modeled two mechanisms through which resistance alleles are generated, *de novo* mutations of wild type alleles, and non-homologous end joining (NHEJ) events during gene drive conversion. In the *de novo* mutation case, the probability that a resistance allele will appear is correlated with the number of wild type alleles in both populations, because only wild type alleles can be converted to resistance alleles. In NHEJ-generated resistance alleles, this probability is correlated with the number of heterozygotes, because NHEJ events occur during conversion, which occurs only in heterozygotes. The spread of the resistance allele, once it has appeared, depends on the presence of the gene drive allele, because otherwise the resistance allele has no fitness advantage over the wild type allele. Therefore, when considering the dynamics in the gene swamping outcome (Fig. 1D), we expect that the duration of the suppression phase with different parameters (Fig. 4B) will affect the probability of resistance evolving.

In Figure 5 we compare the probabilities that resistance does not evolve in the gene swamping and suppression outcomes. These probabilities were computed using stochastic simulations, where we simulated resistance alleles generated by two different mechanisms, following the equations described in [19] (see *Methods*). We considered the robustness of an outcome to the evolution of resistance as the proportion of stochastic simulations leading to the specified outcome in which resistance did not evolve. Since we were interested in the long-term effects of resistance evolution, we considered resistance to have evolved in simulations in which the resistance allele frequency was *≥* 0.1 at the end of the simulation.

**Figure 5:**
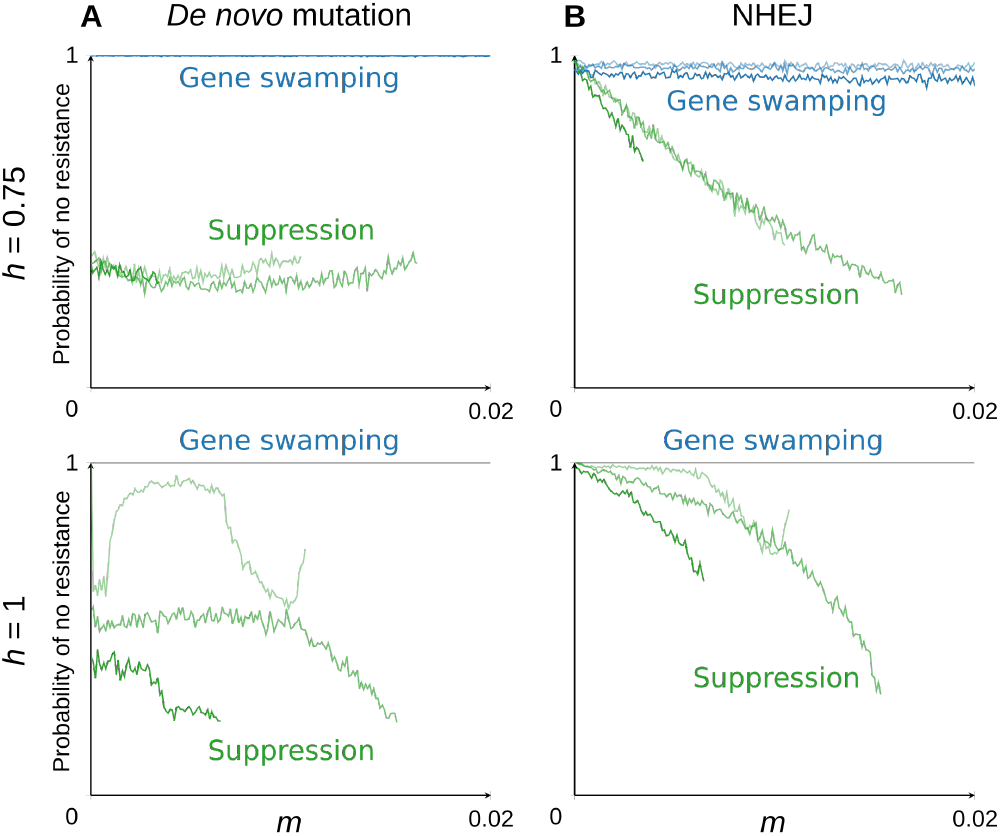
Evolution of resistance in gene swamping and suppression outcomes. Resistance alleles are generated either by *de novo* mutation (A) or non-homologous end joining (NHEJ) during conversion (B). Plotted are the proportions of simulations of gene drive designs leading to ‘gene swamping’ (blue) or ‘suppression’ (green) in which resistance did not evolve, for increasing migration rates. Each line represents a specific gene drive design. Blue lines represent values of *s* that lead to gene swamping outcomes: *s* = 0.85, *s* = 0.8 and *s* = 0.75 for dark, medium and light blue, respectively, in the upper panels (*h* = 0.75), and *s* = 0.7, *s* = 0.65 and *s* = 0.6 for dark, medium and light blue, respectively, in the lower panels (*h* = 1). Green lines represent values of *s* that lead to suppression outcomes: *s* = 0.7, *s* = 0.69 and *s* = 0.68 for dark, medium and light green, respectively, in the upper panels (*h* = 0.75), and *s* = 0.54, *s* = 0.53 and *s* = 0.52 for dark, medium and light green, respectively, in the lower panels (*h* = 1). Other parameters used in all panels: *d* = 10, *c* = 1, *q*_1_(0) = 0.8, *I*_*µ*_ = 0.01, and *I*_*δ*_ = 0.1.

For resistance alleles generated through *de novo* mutations, we observe that gene swamping is more robust than suppression (i.e., fewer simulations resulted in significant spread of a resistance allele), and in both outcomes the probability for resistance evolution does not depend on the migration rate (Fig. 5A). With *h* = 0.5, the evolution of resistance from gene swamping outcomes is more unlikely than with *h* = 0, and did not occur in almost all of our simulations (Fig. 5A). In contrast, for resistance alleles generated through NHEJ events, the difference between the two outcomes is migration-dependent: at low levels of migration, both outcomes have a similar low likelihood of the evolution of resistance, while at higher levels of migration gene swamping is more robust to the evolution of resistance than suppression (Fig. 5B).

The difference between the resistance-allele generating mechanisms, in terms of the effect of migration, relates to the conditions which increase the likelihood of resistance alleles appearing. For *de novo* mutations, the rate of emergence of resistance alleles is a function of the number of wild-type alleles in the system (Eq. 10). Therefore, the suppression of the target population is not sufficient to prevent the evolution of resistance, because *de novo* mutations can appear in the non-target population and spread to the target population through gene flow. For NHEJ events, emergence of resistance is a function of the number of heterozygotes in the system (Eq. 11). For suppression outcomes, higher gene flow results in more intermediate levels of *q* in both populations (i.e. with higher *m, q*_1_ is decreased and *q*_2_ is increased), and therefore a higher proportion of heterozygotes in the system. In gene swamping outcomes, higher gene flow likewise increases the maximal spillover frequency, increasing the frequency of heterozygotes during the suppression phase. This increases the likelihood of resistance evolving in both suppression and gene swamping outcomes, but particularly for suppression due to the persistence of the gene drive allele in the system.

## 4 Discussion

Gene drives have the potential to revolutionize biocontrol, but developing safe deployment approaches is crucial to avoiding unintentional spillover of the gene drive to non-target populations. Feedback between the evolutionary spread of the gene drive and its demographic effects may lead to non-optimal deployment outcomes, such as persistence of the gene drive, while preventing elimination of the target population [8, 20].

Here, we show that this feedback can potentially result in gene drives that avoid spillover and persistence of the drive allele, while still generating strong short-term suppression of the target population without suppression of the non-target population (the ‘gene swamping’ outcome).

Previous models of gene drive spread have taken either a purely soft selection approach [6, 14], in which the gene drive and wild type alleles compete directly, or a hard selection approach, in which the demographic effect of the gene drive determines selection of the gene drive allele [8, 17, 20]. Our model combines these approaches, allowing us to analyze the effect of demographic feedback on deployment outcomes. This feedback, under some conditions, can generate outcomes of gene drive deployment that are absent when considering only evolutionary dynamics (Fig. 1). Notably, the outcomes ‘oscillations’ and ‘gene swamping’ are present only when evolutionary and demographic changes occur in the same timescale (see Supporting Information for a two-timescale model in which these outcomes do not emerge).

One of the main challenges in gene drive deployment is the containment of the released drive to a specific population. Certain gene drive designs are expected to result in uncontrolled spread of the gene drive even at low migration rates [5] (e.g., Fig. 1A), whereas only a limited set of gene drive designs allow differential targeting of the gene drive, at least for low migration rates [6] (Fig. 1B–C). Our results, as well as results from continuous-space and reaction-diffusion models [8, 17], indicate that, when targeted suppression is attempted in the presence of migration, local collapse of a target population or region results in either temporary or persistent suppression, and not in elimination of the populations. In such cases, demographic (Fig. 1C) or spatial [20] oscillations resulting from demographic rescue or gene swamping are a possibility.

We focused our detailed analysis on the ‘gene swamping’ outcome because its characteristics suggest that it could be used as a fail-safe for gene drive deployment. In this scenario, a temporary phase of suppression of the target population is followed by removal of the gene drive from both populations due to wild-type-abundant gene flow from the non-target population (Fig. 1F). This outcome has several interesting features.

(i) In terms of the range of gene drive designs and migration rates leading to this outcome, in comparison to long-term suppression of the target population, the gene swamping outcome is more robust to spillover (Fig. 3A). (ii) The gene swamping outcome is also more robust to the evolution of resistance (Fig. 5). Short-term suppression decreases the chance of resistance evolving during deployment, whereas long-term persistence of the gene drive in the wild increases the probability that resistance to the gene drive will evolve. This may affect the potential for future deployment programs in the same population. (iii) In the gene swamping outcome, at the end of the process the gene drive allele is absent from both populations, but unlike in the ‘failure’ outcome, a targeted suppression phase does occur. While this outcome offers only a confined window of suppression (Fig. 4), non-genetic means of population suppression could be used during this window to achieve eradication of the target population. However, as opposed to long-term suppression, in the case that eradication is not achieved, the deployment will not result in spillover nor in the gene drive allele remaining in the system. Since these non-genetic control measures should be timed to coincide with the decrease in the population size of the target population, control measures that are efficient at low population densities, such as mating disruption strategies (e.g., false pheromones), should be preferred over measures such as pesticides or sterile males [27].

The investigation of gene drive deployment strategies has progressed mainly using mathematical and computational modeling. However, for the conclusions from these modeling efforts to be incorporated into upcoming gene drive projects, it is likely that experimental validation in a lab setting would be required. So far, experimental setups for population suppression using gene drives have been conducted using single caged populations [28] (usually with several replicates). In order to demonstrate the potential for spillover mitigation with different gene drive configurations, it would be important to design an experimental setup that includes population structure, and that strongly relates to the setup of theoretical models. In this aspect, the modeling approach of two interconnected populations, as we have used here, is perhaps more readily translatable to an experimental setup, for example using two interconnected cages, compared to continuous-space models.

Our model demonstrates the importance of considering the interaction between the evolutionary and demographic effects of gene drives. With such feedback, qualitatively different outcomes may emerge, which may have implications for designing gene drive configurations and deployment strategies. In our model, asymmetrical gene flow between target and non-target populations was shown to generate an outcome of interest which we termed ‘gene swamping’. This fail-safe strategy does not rely on the complexity of the gene drive construct [29–31], but rather on gene flow and population dynamics. Since these aspects are general features of wild populations, this approach could be readily combined with other types of fail-safe mechanisms to prevent the unintended spread of gene drive constructs.

## 5 Data Accessibility

To facilitate further exploration of our results, we uploaded the model to the platform `modelRxiv`, which allows visualization and analysis of models through a simple user interface ([21]; https://modelrxiv.org/model/yoKkSv and https://modelrxiv.org/model/k8h5M6). In this platform, the results can be regenerated and the model can be re-parameterized (Fig. S9). The model code is available on modelRxiv at the URLs noted above, GitHub (https://github.com/carrowkeel/genedrive_demographic) and Zenodo (https://doi.org/10.5281/zenodo.7371644).

## 6 Acknowledgements

We would like to thank Adam Lampert for insightful comments. This project was supported by Israel Science Foundation (ISF) Grant 2049/21, and by German-Israel Foundation (GIF) Grant I-1526-500.15/2021.

## 7 Author contributions

KDH and GG developed the model. KDH analyzed the model and generated figures. KDH and GG co-authored the manuscript. GG supervised and acquired funding for the research.

## 9 Methods

### 9.1 Population dynamics

The population dynamics are governed by Eqs. 1 and 4. In Eq. 4, when *Ñ* = *K* the population size would remain fixed from that point onward. To avoid the population size remaining fixed at *K* when we initialize the dynamics, we defined the initial population size of both populations as *N*_*i*_(0) = *K −EK*. We used *E* = 0.01 through all analyses. Changing the value of *E* changes the delay of the demographic effect of the gene drive: lower values delay the demographic effect, while higher values increase the rate of the demographic effect.

### 9.2 Classification of outcomes

We classified the outcomes based on the evolutionary (Eq. 3) and the demographic (Eq. 4) dynamics. To differentiate between outcomes, we consider (i) the gene drive allele frequency and population size at *t* = 100, and (ii) the minimum population size. For each outcome, we define thresholds of these two parameters. For example, we classify the outcome as ‘suppression’ when the target population has a high frequency (*≥* 0.5) of the gene drive allele, the non-target population has a low frequency (*<* 0.5) of the gene drive allele, the target population is suppressed (*N*_1_ *<* 0.9*K*, i.e., less than 90% of its initial size), and the non-target population was not suppressed (min(*N*_2_) *≥* 0.9*K*). The full classification of outcomes is detailed in Table S1.

We defined three characteristics of the suppression phase in the target population (Fig. 4A): (i) the duration of suppression, (ii) the maximal suppression level, and (iii) the total suppression. The duration of suppression was defined as the number of generations in which the target population size is suppressed (*N*_1_ *<* 0.9*K*, i.e., less than 90% of its initial size). The maximal suppression level is defined as 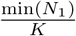. The total suppression is defined as 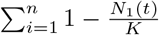 where *n* is the total number of generations.

### 9.3 Evolution of resistance

In order to measure the effect of the evolution of resistance on the outcomes of our model, we modified the genetic element of the model, using a previous model [19] which considers three possible alleles at the gene drive locus: *a* (wild type allele), *A* (gene drive allele) and *r* (resistance allele). Here, in addition to the gene drive allele, we track the frequency of the resistance allele in the two populations, *p*_*i*_. To account for gene flow of the resistance allele, we add an equation analogous to Eq. 2 to compute the post-migration resistance allele frequency 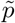:

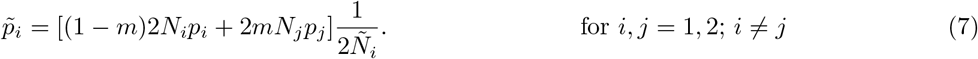

Because the resistance allele prevents conversion, we only need to account for conversion for *A/a* heterozygotes. We define *s*_*r*_ as the fitness cost of the resistance allele, and *h*_*r*_ as the degree of dominance of the resistance allele. Using these parameters, we define the evolutionary dynamics in a similar manner as in Eq. 3 for each population following [19]:

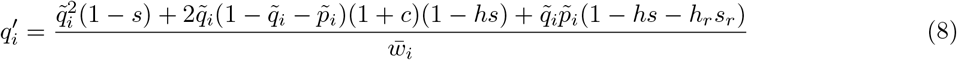

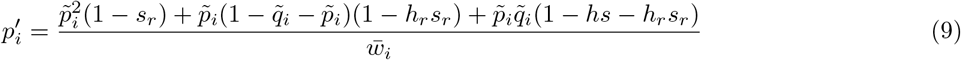

These recurrence equations incorporate the gene drive design parameters (*s, c* and *h*) and the fitness cost and dominance of the resistance allele (*s*_*r*_ and *h*_*r*_, respectively). Accounting for the resistance allele *p*_*i*_, the mean fitness in the two populations is now 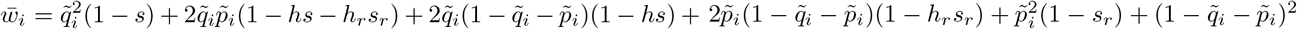 For the purpose of investigating the likelihood of resistance establishing in the population we are mainly interested in the emergence of a significant level of resistance alleles in either population, and not in the loss of resistance alleles following the loss of the gene drive allele (through competition with the wild type allele). Therefore, we set the relative fitness cost of the resistance allele to *s*_*r*_ = 0. This implies that, in our analysis, the frequency of resistance alleles will remain at a fixed level once the gene drive allele is lost because it has no disadvantage relative to the wild type allele. This assumption is useful for the evaluation of the maximal spread of the resistance allele under different conditions.

We considered two mechanisms through which resistance alleles are generated [19]: *de novo* mutations of the wild type allele to a resistance allele that cannot be converted, and non-homologous end joining during conversion resulting in a resistance allele (as opposed to a converted gene drive allele). *De novo* mutations appear as a function of population size and wild type alleles [19]:

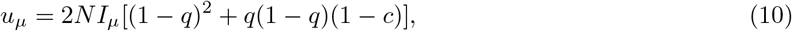

where *u*_*µ*_ is the probability of a *de novo* mutation appearing in the current generation, *N* is the relative population size (in relation to the carrying capacity *K*), and 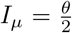 is the scaled *de novo* mutation rate in a population of size *K* (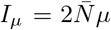, where *µ* is the mutation rate and 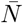 is the effective population size at carrying capacity). For our simulations, we define *I*_*µ*_ = 0.01. (1 *− q*)^2^ and *q*(1 *− q*)(1 *− c*) are the proportions of wild type homozygotes and half the non-converted heterozygotes, respectively.

In the case of NHEJ, events occur as a function of population size and gene drive/wild type heterozygotes [19]:

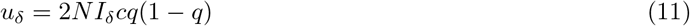

where *u*_*δ*_ is the probability of an NHEJ event occurring in the current generation, *N* is the relative population size, and 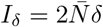 is the rate of NHEJ events per generation in a population of size *K*. For our simulations, we define *I*_*δ*_ = 0.1. *cq*(1 *− q*) is the proportion of converted heterozygotes.

To compute the probability of resistance evolving either through *de novo* mutation or NHEJ events, we defined the robustness of an outcome to the evolution of resistance as the proportion of simulations in which resistance did not evolve. For each simulation run, we defined the evolution of resistance according to the resistance allele frequency *p* at the end of the simulation, where *p >* 0.1 was considered as resistance having evolved. We ran 1000 simulation repeats for different gene drive designs (different values of *s*) for recessive (*h* = 0) and additive (*h* = 0.5) gene drive alleles within a range of migration rates *m*.

## References

1. Alphey, L. S., Crisanti, A., Randazzo, F. F. & Akbari, O. S. Opinion: Standardizing the definition of gene drive. Proceedings of the National Academy of Sciences 117, 30864–30867 (2020).

2. Gantz, V. M. et al. Highly efficient Cas9-mediated gene drive for population modification of the malaria vector mosquito Anopheles stephensi. Proceedings of the National Academy of Sciences 112, E6736– E6743 (2015).

3. Hammond, A. et al. A CRISPR-Cas9 gene drive system targeting female reproduction in the malaria mosquito vector Anopheles gambiae. Nature Biotechnology 34, 78–83 (2016).

4. Bier, E. Gene drives gaining speed. Nature Reviews Genetics 23, 5–22 (2022).

5. Noble, C., Adlam, B., Church, G. M., Esvelt, K. M. & Nowak, M. A. Current CRISPR gene drive systems are likely to be highly invasive in wild populations. eLife 7, e33423 (2018).

6. Greenbaum, G., Feldman, M. W., Rosenberg, N. A. & Kim, J. Designing gene drives to limit spillover to non-target populations. PLoS Genetics 17, e1009278 (2021).

7. Price, T. A. et al. Resistance to natural and synthetic gene drive systems. Journal of Evolutionary Biology 33, 1345–1360 (2020).

8. Girardin, L. & Débarre, F. Demographic feedbacks can hamper the spatial spread of a gene drive. Journal of Mathematical Biology 83, 67 (2021).

9. James, S. et al. Pathway to deployment of gene drive mosquitoes as a potential biocontrol tool for elimination of malaria in sub-Saharan Africa: recommendations of a scientific working group. The American Journal of Tropical Medicine and Hygiene 98, 1–49 (2018).

10. James, S. L., Marshall, J. M., Christophides, G. K., Okumu, F. O. & Nolan, T. Toward the definition of efficacy and safety criteria for advancing gene drive-modified mosquitoes to field testing. Vector-Borne and Zoonotic Diseases 20, 237–251 (2020).

11. Hartl, D. Analysis of a general population genetic model of meiotic drive. Evolution 24, 538–545 (1970).

12. Burt, A. Site-specific selfish genes as tools for the control and genetic engineering of natural populations. Proceedings of the Royal Society of London. Series B: Biological Sciences 270, 921–928 (2003).

13. Deredec, A., Burt, A. & Godfray, H. C. J. The population genetics of using homing endonuclease genes in vector and pest management. Genetics 179, 2013–2026 (2008).

14. Unckless, R. L., Messer, P. W., Connallon, T. & Clark, A. G. Modeling the manipulation of natural populations by the mutagenic chain reaction. Genetics 201, 425–431 (2015).

15. Verma, P., Reeves, R. G. & Gokhale, C. S. A common gene drive language eases regulatory process and eco-evolutionary extensions. BMC Ecology and Evolution 21, 156 (2021).

16. Tanaka, H., Stone, H. A. & Nelson, D. R. Spatial gene drives and pushed genetic waves. Proceedings of the National Academy of Sciences 114, 8452–8457 (2017).

17. Champer, J., Zhao, J., Champer, S. E., Liu, J. & Messer, P. W. Population dynamics of underdominance gene drive systems in continuous space. ACS Synthetic Biology 9, 779–792 (2020).

18. Gomulkiewicz, R., Thies, M. L. & Bull, J. J. Evading resistance to gene drives. Genetics 217, iyaa040 (2021).

19. Unckless, R. L., Clark, A. G. & Messer, P. W. Evolution of resistance against CRISPR/Cas9 gene drive. Genetics 205, 827–841 (2017).

20. Champer, J., Kim, I. K., Champer, S. E., Clark, A. G. & Messer, P. W. Suppression gene drive in continuous space can result in unstable persistence of both drive and wild-type alleles. Molecular Ecology 30, 1086–1101 (2021).

21. Harris, K. D., Hadari, G. & Greenbaum, G. modelRxiv: A platform for the distribution, computation and interactive display of models. bioRxiv. eprint: https://www.biorxiv.org/content/early/2022/02/19/2022.02.16.480599.full.pdf (2022).

22. Haldane, J. B. S. The relation between density regulation and natural selection. Proceedings of the Royal Society of London. Series B-Biological Sciences 145, 306–308 (1956).

23. Thomé, R. C., Yang, H. M. & Esteva, L. Optimal control of Aedes aegypti mosquitoes by the sterile insect technique and insecticide. Mathematical Biosciences 223, 12–23 (2010).

24. Marshall, J. M., Buchman, A., Sánchez C, H. M. & Akbari, O. S. Overcoming evolved resistance to population-suppressing homing-based gene drives. Scientific reports 7, 1–12 (2017).

25. Prowse, T. A. et al. Dodging silver bullets: good CRISPR gene-drive design is critical for eradicating exotic vertebrates. Proceedings of the Royal Society B: Biological Sciences 284, 20170799 (2017).

26. Champer, J. et al. Novel CRISPR/Cas9 gene drive constructs reveal insights into mechanisms of resistance allele formation and drive efficiency in genetically diverse populations. PLoS genetics 13, e1006796 (2017).

27. Lampert, A. & Liebhold, A. M. Combining multiple tactics over time for cost-effective eradication of invading insect populations. Ecology Letters 24, 279–287 (2021).

28. Hammond, A. et al. Gene-drive suppression of mosquito populations in large cages as a bridge between lab and field. Nature Communications 12, 1–9 (2021).

29. Noble, C. et al. Daisy-chain gene drives for the alteration of local populations. Proceedings of the National Academy of Sciences 116, 8275–8282 (2019).

30. Li, M. et al. Development of a confinable gene drive system in the human disease vector Aedes aegypti. eLife 9, e51701 (2020).

31. Xu, X.-R. S. et al. Active genetic neutralizing elements for halting or deleting gene drives. Molecular Cell 80, 246–262 (2020).

