## Supplemental Information 1 for "Rescue by gene swamping as a gene drive deployment strategy"

### Supplementary Material: Rescue by gene swamping as a gene drive deployment strategy

#### Contents


**1 Modeling the relationship between selection and demography** **2**

1.1 Effect of $r_1$ formulation on deployment outcomes . . . . . . . . . . . . . . . . . . . . . . . . . 4

**2 Definition of deployment outcomes** **5**

**3 The effect of the initial frequency of the gene drive on deployment outcomes** **6**

**4 Characterizing suppression in gene swamping outcomes** **7**

**5 Modeling with separation of timescales** **8**

**6 Zygotic conversion** **10**

**7 Exploration of model parameter space using `modelRxiv`** **12**

### 1 Modeling the relationship between selection and demography

In order to define the relationship between the relative fitness cost of the gene drive $s$ and the growth rate of a population with a gene drive frequency $q$, we introduced three different growth rate parameters: $r_0$, the growth rate of a population of homozygotes of the wild type allele; $r_h$, the growth rate of a population of heterozygotes; and $r_1$, the growth rate of a population of homozygotes of the gene drive allele. We defined the growth rate of a population ($R$) as the sum of these growth rates weighted by the proportion of the respective genotypes, assuming Hardy-Weinberg equilibrium frequencies (Eq. 5 in the main text). We defined $r_h$ as the weighted sum of $r_0$ and $r_1$ using the dominance parameter of the gene drive allele $h$.

To allow flexibility in the degree to which demographic feedback is possible, we introduced a parameter $d$ that determines the relationship between $s$ and $r_1$ (Eq. 6 in the main text). At $d = 0$, there is no demographic effect, i.e., $r_1 = r_0$. When $d \to \infty$, $r_1 \to -s$. In other words, at high values of $d$, the effect of the gene drive on selection is correlated to its effect on growth rate (Fig. S1A). It is important to note that for the population $i$ to decrease in size, we need $R_i < 0$.

Using a parameter to determine the effect of the gene drive on demography allows us to observe how demographic feedback changes the deployment outcome, across a continuum from purely soft selection ($d = 0$), to a full correlation between selection and demography ($d \to \infty$). Increasing $d$, we observe a shift in the deployment outcome (Fig. S1B). Specifically, there is a shift from 'targeted outcomes', in which the gene drive and demographic effects are targeted to one population ('differential targeting', 'suppression', 'oscillations' and 'gene swamping') to 'non-targeted outcomes', in which there is no targeting ('spillover' and 'collapse'). The dashed line in Figure S1B indicates a threshold of $m$ below which targeted outcomes are achievable; increasing $d$ increases the threshold of $m$, allowing targeted outcomes at higher migration rates for the same gene drive design.

We can expand the above analysis to a wider range of gene drive designs by defining a threshold value $m^*$ below which targeted outcomes exist for some initial frequency of the gene drive. For this analysis we define $q_1(0) = 1, q_2(0) = 0$, and compute the maximal value of $m$ that gives a targeted outcome, for different gene drive configurations ($c = 1$, with different values of $s$). This definition of $m^*$ is a numerically-computed equivalent of the definition of $m^*$ in [1]. As shown in Figure S1B, increasing $d$ increases the migration threshold for targeted outcomes, but this effect depends on the gene drive design (Fig. S1C–D).

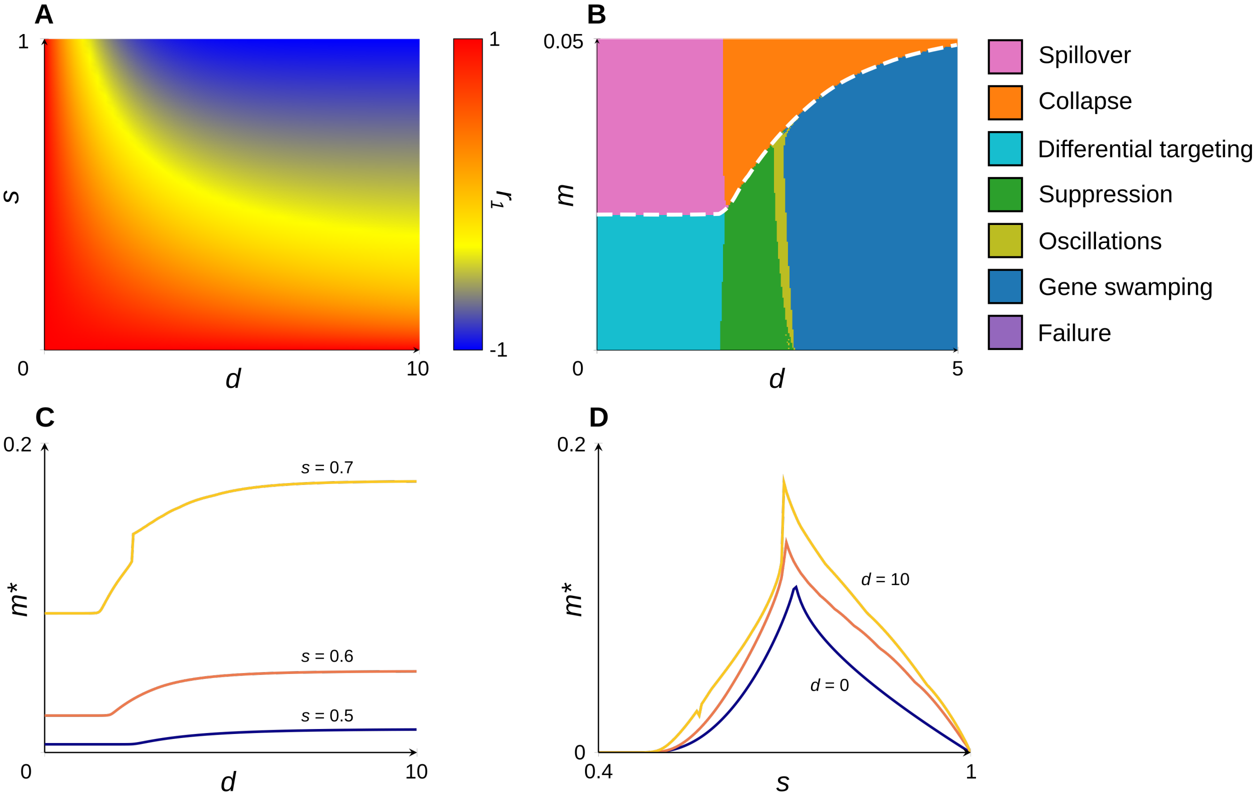


Figure S1: **Effect of demographic parameter ($d$) on outcomes.** (A) Relationship between the demographic effect of the gene drive (the growth rate $r_1$) and the fitness cost $s$ of the gene drive, as a function of $d$. (B) Change in deployment outcome for a specific gene drive design as a function of migration ($m$) and $d$. Colors relate to outcomes as indicated by the legend. The white dashed line indicates a value of $m$ below which targeted outcomes are achievable. Other parameters used are $s = 0.6$, $c = 1$ and $q_1 = 0.8$. (C) Migration threshold ($m^*$) for targeted outcomes ('differential targeting', 'suppression', 'oscillations' and 'gene swamping'), for different levels of demographic feedback ($d$) and relative fitness costs ($s$). (D) Migration threshold for targeted outcomes of gene drive designs for different levels of demographic feedback ($d$). For (C–D), other parameters used are $c = 1$ and $q_1 = 1$.

#### 1.1 Effect of $r_1$ formulation on deployment outcomes

In our model, the definition of the growth rate of a population fixed for the gene drive ($r_1$) provides the link between the evolutionary and demographic dynamics. In the main model we used a formulation of this relationship that provided a continuum between purely evolutionary dynamics ($r_1 = r_0$), and a full correlation between the fitness cost of the gene drive $s$ and its effect on growth rate (Eq. 6 in the main text). To demonstrate that different formulations of $r_1$ give a qualitatively similar result in terms of deployment outcomes, we analyzed an alternative formulation of $r_1$ alongside the formulation used for all other analyses (Eq. 6 in the main text). This formulation includes the fitness cost of the gene drive allele $s$ and the parameter $d$, but lacks $r_0$:

$$r_1 = -ads, \tag{S1}$$

where $a$ is a scaling parameter. The proportions of outcomes in which either or both target and non-target populations are suppressed ('collapse', 'suppression', 'oscillations' and 'gene swamping') increase with higher values of $d$ for both formulations (Fig. S2). While there is a difference between the formulations in the proportion of each outcome for the same level of migration, the behavior of each outcome under increasing migration rates is similar between the formulations (Fig. S2). In other words, the formulation of $r_1$ does not change the types of outcomes produced by the model, or alter the relationship between these outcomes qualitatively.

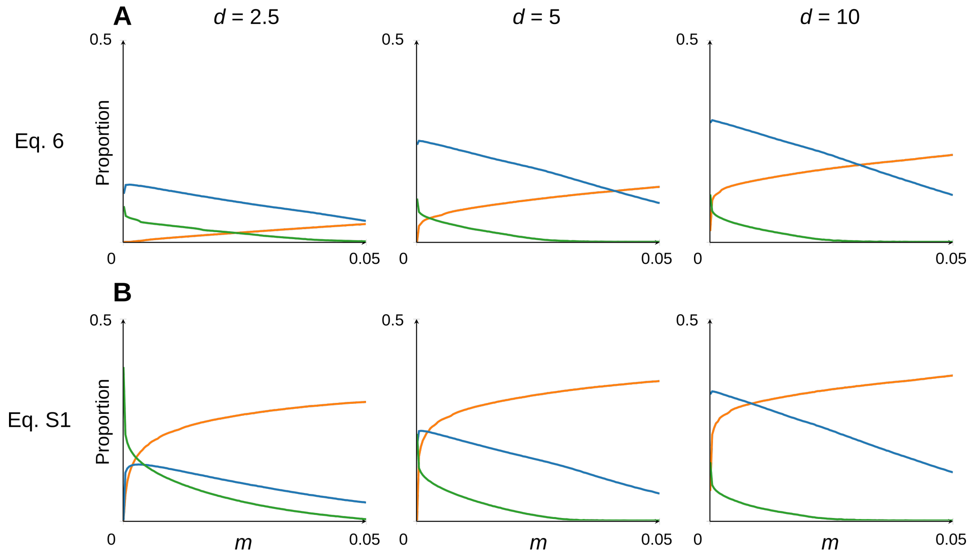


Figure S2: **Effect of the formulation of $r_1$ on the proportion of deployment outcomes**. (A) The proportions of gene drive designs leading to collapse (orange), suppression or oscillations (green) and gene swamping (blue) outcomes at different levels of demographic effect ($d$) for increasing migration rates ($m$), for the formulation of $r_1$ used in the main text (Eq. 6). (B) The proportions of gene drive designs leading to collapse (orange), suppression or oscillations (green) and gene swamping (blue) outcomes at different levels of demographic effect ($d$) for increasing migration rates ($m$), for a simplified formulation of $r_1$ (Eq. S1). Other parameters used are $h = 1$, $q_1(0) = 0.8$, and $a = 0.1$ (the scaling parameter for $d$).

#### 2 Definition of deployment outcomes

To differentiate between deployment outcomes, we defined criteria for classifying model dynamics ($q_i$ and $N_i$) (see *Methods* section). Outcome definitions take into account, for both populations, the frequency of the gene drive at $t = 100$, and population size at $t = 100$ as well as the minimum population size throughout the 100 generations of the simulation. These definitions allows us to connect outcomes to functional aspects of deployment (e.g., the minimum population size is a measure of the effectiveness of suppression during deployment). For all outcomes except 'oscillations', the outcome was defined according to thresholds of these parameters (Table S1). For 'oscillations', in addition to the thresholds in this table, we consider dynamics in which the target population size oscillates with an amplitude that does not decrease over time.

| **Outcome** | $q_1(n)$ | $q_2(n)$ | $N_1(n)$ | $N_2(n)$ | $\min\{N_1\}$ | $\min\{N_2\}$ |
|---|---|---|---|---|---|---|
| Spillover | $> 0.5$ | $> 0.5$ | $\geq 0.9$ | $\geq 0.9$ | $\geq 0.9$ | $\geq 0.9$ |
| Collapse | $> 0.5$ | $> 0.5$ | $< 0.9$ | $< 0.9$ | $< 0.9$ | $< 0.9$ |
| Failure | $< 0.5$ | $< 0.5$ | $\geq 0.9$ | $\geq 0.9$ | $\geq 0.9$ | $\geq 0.9$ |
| Differential targeting, no collapse | $> 0.5$ | $< 0.5$ | $\geq 0.9$ | $\geq 0.9$ | $\geq 0.9$ | $\geq 0.9$ |
| Gene swamping | $< 0.5$ | $< 0.5$ | $\geq 0.9$ | $\geq 0.9$ | $< 0.9$ | $\geq 0.9$ |
| Oscillations | $> 0$ | $< 0.5$ | * | $\geq 0.9$ | $< 0.9$ | $\geq 0.9$ |
| Suppression | $> 0.5$ | $< 0.5$ | $< 0.9$ | $\geq 0.9$ | $< 0.9$ | $\geq 0.9$ |

Table S1: **Definition of outcomes**. Outcomes are classified according to the thresholds denoted in this table. The 'oscillations' outcome additionally requires that the target population size oscillates with an amplitude that does not decrease over time.

#### 3 The effect of the initial frequency of the gene drive on deployment outcomes

In all analyses of outcomes, we used a relatively high initial frequency of the gene drive in the target population ($q_1(0) = 0.8$). This means deploying four times the population size in gene drive homozygotes. Because this scenario may not be achievable in some systems, we analyzed outcomes for a range of values of $q_1$, equivalent to deployment from one to fives times the population size in gene drive homozygotes. While at lower values of $q_1(0)$ targeted suppression exists for a narrower range of gene drive designs, both suppression and gene swamping outcomes are still achievable (Fig. S3).

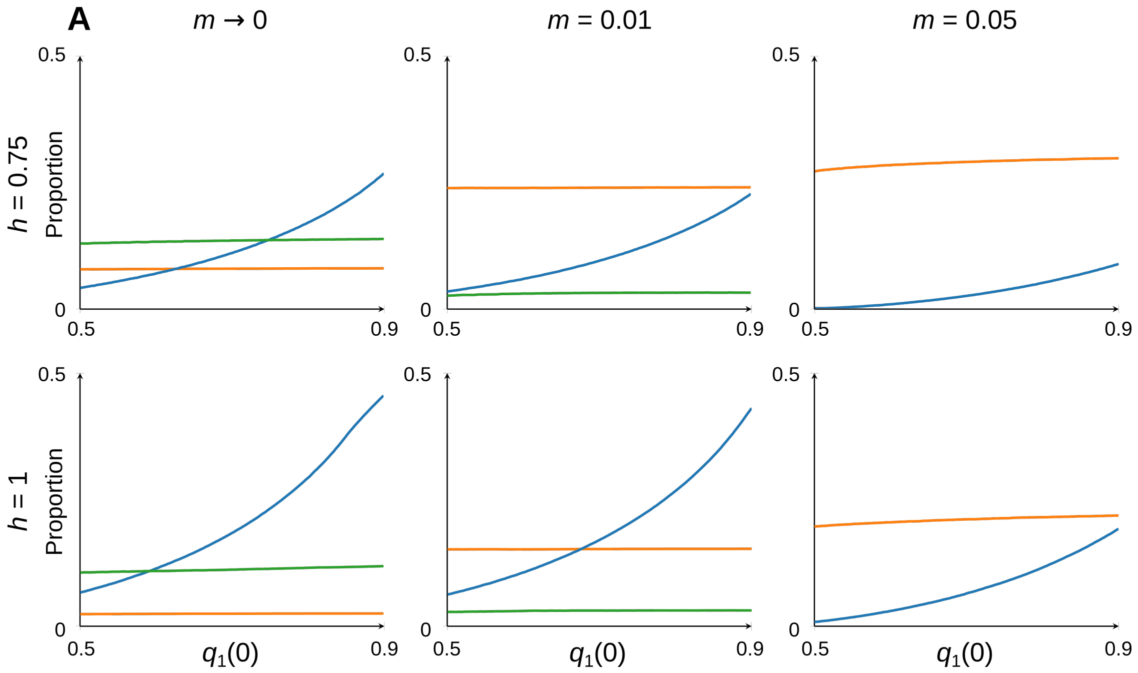


Figure S3: **Effect of initial frequency and demography on outcome proportions.** Corresponding to Figure 2 in main text. (A) Proportions of gene drive designs (i.e., values of $s$, $c$ and $h$) leading to collapse (orange), suppression (green) and gene swamping (blue) outcomes at different rates of migration, for increasing values of the initial frequency of the gene drive in the target population ($q_1(0)$). Other parameters used are $d = 10$ and $q_2(0) = 0$.

### 4 Characterizing suppression in gene swamping outcomes

In the 'gene swamping' outcome, the target population undergoes short-term suppression followed by loss of the gene drive from both populations. In order to understand how the suppression phase in gene swamping is affected by gene drive design and migration, we studied three aspects of suppression (Fig. 4A): (i) the number of generations in which the target population is beneath the suppression threshold we defined (i.e., $N_1 < 0.9K$) ('duration'), (ii) the maximal suppression level of the target population ('suppression level'), and (iii) the accumulative suppression of the target population ('total suppression'). In the main text we investigated these three aspects for different values of $s$ and for $m = 0.01$; here we investigate in addition the effect of different migration rates (Fig. S4). We found that the suppression phase is longer, and the suppression level higher for lower values of $s$; increased levels of migration affect mainly the duration of suppression, but also reduce the suppression level (Fig. S4). Higher migration increases the rate of gene swamping, shortening the suppression phase.

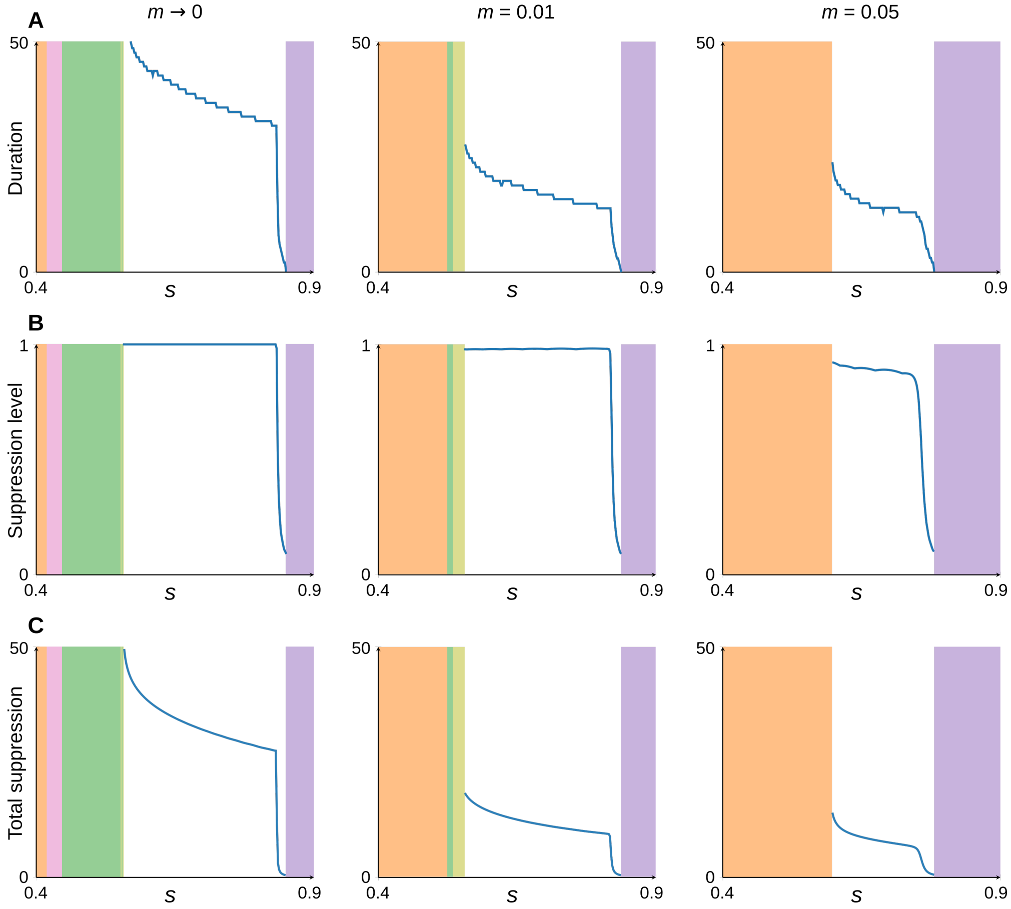


Figure S4: **Effect of gene drive design and migration on suppression characteristics in gene swamping.** Corresponding to Figure 4 in main text. Blue curves indicate the value of each parameter for gene swamping outcomes; values of $s$ leading to other outcomes are colored according to the color legends in Figure S1. (A-C) Behavior of different suppression characteristics for a range of different gene drives (values of $s$), for different migration rates ((A) Duration, (B) Suppression level, and (C) Total suppression). Other parameters used are: $d = 10$, $c = 1$, and $q_1(0) = 0.8$.

#### 5 Modeling with separation of timescales

The model in the main text was designed with the appreciation that in gene drive deployments demographic changes occur in the same timescale as evolutionary changes. This is based on the fact that the fast-spreading, highly deleterious gene drive allele is designed to impact the demographic equilibrium. For the timescales to be separate, the population size must reach a new equilibrium faster than the evolutionary spread of the gene drive.

In order to investigate the importance of the similarity in timescales for ascertainment of the outcomes we observe in the model presented in the main text, we generated and studied a simplified version of our model where the time scales are separated. To do this, we modified Eq. 4 in the main text so that population size directly follows the frequency of the gene drive:

$$N_i' = 1 - (q_i')^{d^*}. \tag{S2}$$

We introduce a parameter $d^*$ to allow control over the demographic effects of the gene drive, as the equivalent of $d$ in the main model. When $d^* = 1$, the population size is linearly correlated to the frequency of the gene drive. For higher values of $d^*$, the effect of the gene drive on population size is reduced. As $N_i'$ is independent of $N_i$, Eq. 1 in the main text is irrelevant for this separate timescale model.

We analyzed the outcomes of the model, using the same criteria as for the full model, to determine the effect of the separation of timescales on potential outcomes, and the relationship between migration and demographic effects on outcome proportions (Figs. S5–S6). We observe that the oscillations of population size that emerged for some parameterizations in the targeted suppression outcomes of the single-timescale model are absent from the two-timescale model. This is because there is a lack of delay between evolutionary and demographic changes in the two-timescales model: the frequency of the gene drive in both populations at $t = n$ directly determines the gene flow at $t = n + 1$. In this model, differential targeting of the gene drive leads to a stable, suppressed population size (Fig. S5). However, demography can still impact the outcome of deployment: for higher values of $d^*$ (that is, the gene drive has a smaller effect on demography), the proportion of suppression outcomes is higher. This is owing to the effect of gene flow from the non-target population to the target population when the latter is suppressed. This demonstrates that the feedback we identified depends, at least in part, on evolutionary and demographic changes occurring at the same timescale.

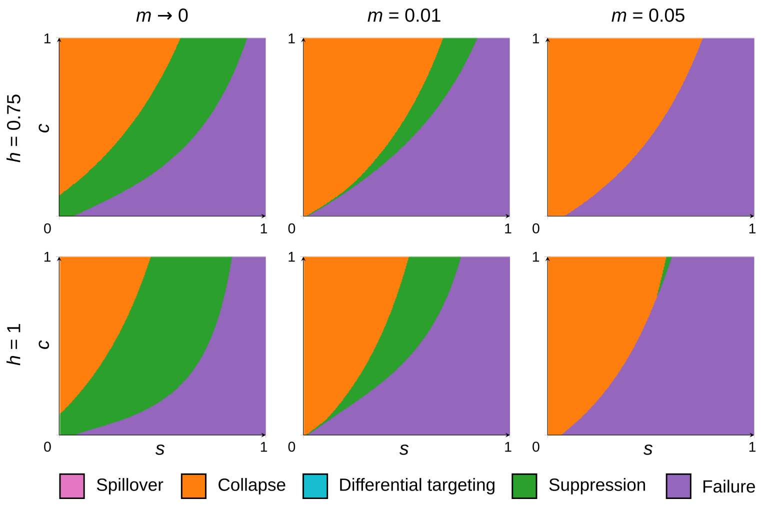


Figure S5: **Two-timescale model outcomes in relation to gene drive design and migration rates ($d^* = 1$).** (A) Each panel shows the outcome attained in our model for a fixed $h$ value and for different $m$ values. In all panels the parameters $d^* = 1$ and $q_1(0) = 0.8$ were used.

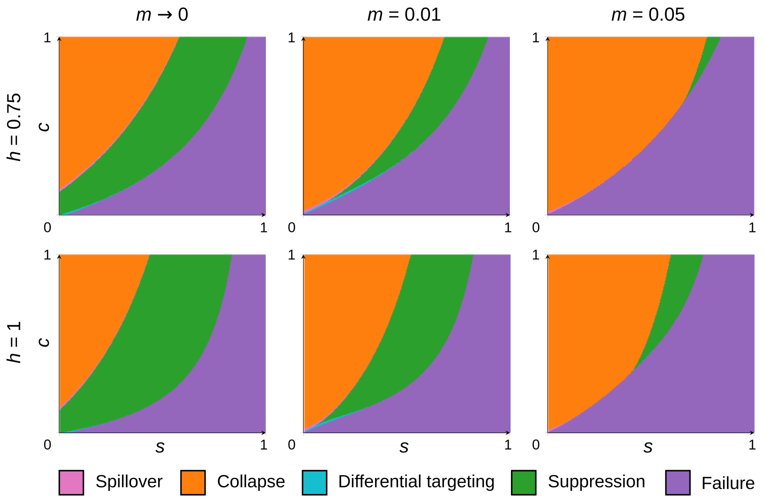


Figure S6: **Two-timescale model outcomes in relation to gene drive design and migration rates ($d^* = 10$).** (A) Each panel shows the outcome attained in our model for a fixed $h$ value and for different $m$ values. In all panels the parameters $d^* = 10$ and $q_1(0) = 0.8$ were used.

#### 6 Zygotic conversion

In the main text we developed a model based that considered gametic conversion, i.e., the CRISPR mechanism is active in the germline, converting heterozygotes to homozygotes. With this mechanism selection that acts on converted heterozygotes is the same as for non-converted heterozygotes, assuming that the gametes themselves are not exposed to selection induced by the gene drive. Another type of conversion mechanism to consider is zygotic conversion [2], where converted heterozygotes undergo the same selection as gene drive homozygotes. While early studies of gene drives assumed that the CRISPR conversion mechanism was active in the zygote [3], current gene drives designs are based on gametic conversion owing to higher conversion efficiency ([4]).

To model conversion in the zygote, we introduce two selection coefficients for converted and non-converted heterozygotes, $s_c$ and $s_n$, respectively, as in [1]. The selection coefficient of converted heterozygotes is defined as $s_c = c(1-s)$, and the selection coefficient of non-converted heterozygotes is defined as $s_n = \frac{1}{2}(1-c)(1-hs)$. We modify Eq. 2 in the main text to incorporate $s_c$ and $s_n$:

$$q_i' = \frac{\tilde{q}_i^2(1-s) + \tilde{q}_i(1-\tilde{q}_i)(s_n + s_c)}{\bar{w}_i}, \tag{S3}$$

where the average fitness $\bar{w}$ is defined as $\bar{w}_i = \tilde{q}_i^2(1-s) + 2\tilde{q}_i(1-\tilde{q}_i)(2s_n + s_c) + (1-\tilde{q}_i)^2$. We also modify Eq. 5 in the main text so that the growth rate of converted heterozygotes is the same as for homozygotes:

$$R_i = (1-q_i)^2 r_0 + 2q_i(1-q_i)[(1-c)r_h + cr_1] + q_i^2 r_1. \tag{S4}$$

For $h = 1$, the zygotic and gametic conversion models converge ([1, 5]). For recessive ($h = 0$) and additive ($h = 0.5$) gene drives, zygotic conversion offers more targeted outcomes than gametic conversion (Figs. S7-S8). This is because gene drive designs with $h \leq 0.5$ are not threshold-dependent with gametic conversion ([5]). While gametic and zygotic conversion models differ quantitatively in terms of outcomes for different parameters for $0 \leq h \leq 1$, the same types of outcomes can be observed under both mechanisms (Fig. S8). In other words, the conversion mechanism does not qualitatively alter the dynamics we observe.

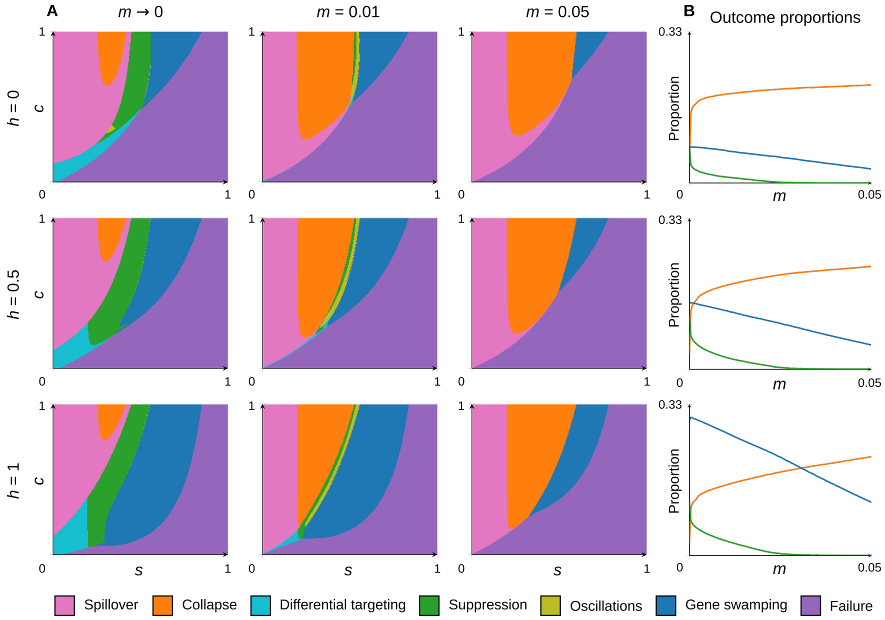


Figure S7: **Zygotic conversion model outcomes in relation to gene drive design and migration rates**. (A) Each panel shows the outcome attained in our model for a fixed $h$ value and for different $m$ values. In all panels the parameters $d = 10$ and $q_1(0) = 0.8$ were used. At $h = 1$, the outcomes converge to those of the gametic conversion model. This figure is comparable to Figure 2 in the main text.

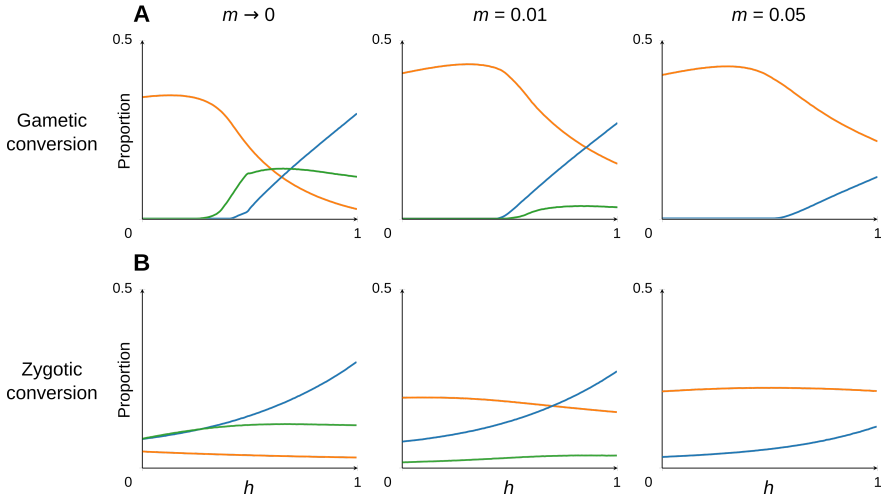


Figure S8: **Effect of gene drive dominance on the outcomes of gene drive deployment assuming gametic or zygotic conversion**. (A-B) Proportions of gene drive designs (i.e., values of $s$, $c$ and $h$) leading to collapse (orange), suppression (green) and gene swamping (blue) outcomes at different rates of migration, for increasing values of dominance of the gene drive allele ($h$), assuming gametic (A) and zygotic (B) conversion. Other parameters used are $d = 10$ and $q_1(0) = 0$.

### 7 Exploration of model parameter space using `modelRxiv`

In order to allow readers to explore the model parameter space beyond the results presented here, we added our model to the `modelRxiv` platform ([6]; `https://modelrxiv.org/model/yoKkSv` and `https://modelrxiv.org/model/k8h5M6`). On the platform, users can manipulate the model parameters and analyze the effect on the model dynamics and outcomes. This can be done either by manually manipulating the model parameters, or by selecting 'presets' that relate to specific figures in the manuscript (Fig. S9). Selecting a preset will modify the selected parameters and run the specific analysis that produced the figure; users can then edit the parameters and rerun the analysis, to understand the effect of these parameters on the result.

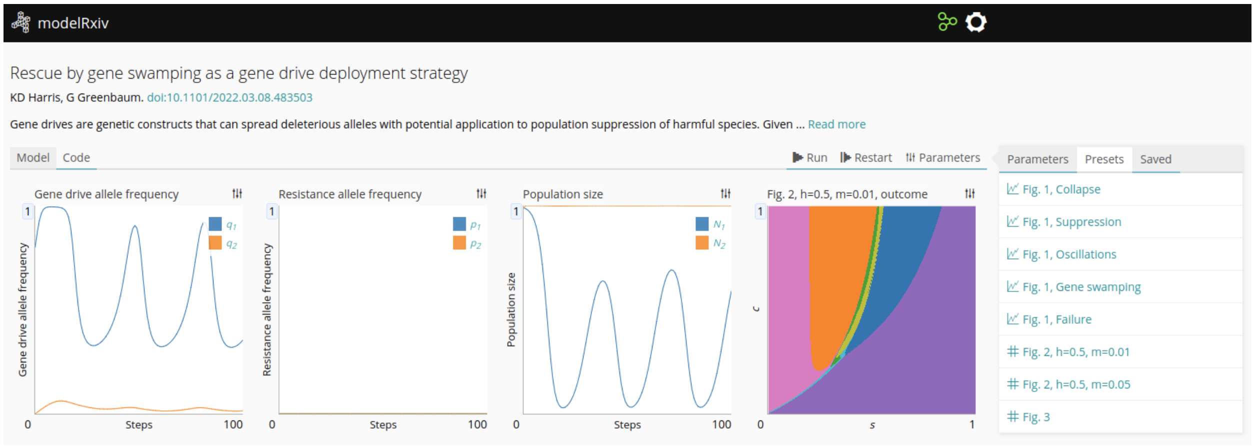


Figure S9: **Model analysis page on `modelRxiv` platform.** Users can visualize dynamics for different parameters using the 'Parameters' menu on the right, or select 'Presets' that relate to figures in the manuscript. After selecting a figure, users can manipulate the model parameters in the 'Parameters' panel and then either run the dynamics of the model by pressing 'Run' or calculate the grid (indicated in the presets menu by a hash symbol beside the preset) by clicking the 'Grid' button.

### References

1. Greenbaum, G., Feldman, M. W., Rosenberg, N. A. & Kim, J. Designing gene drives to limit spillover to non-target populations. *PLoS Genetics* **17,** e1009278 (2021).

2. Grunwald, H. A. *et al.* Super-Mendelian inheritance mediated by CRISPR–Cas9 in the female mouse germline. *Nature* **566,** 105–109 (2019).

3. Gantz, V. M. *et al.* Highly efficient Cas9-mediated gene drive for population modification of the malaria vector mosquito *Anopheles stephensi. Proceedings of the National Academy of Sciences* **112,** E6736–E6743 (2015).

4. Champer, J. *et al.* Novel CRISPR/Cas9 gene drive constructs reveal insights into mechanisms of resistance allele formation and drive efficiency in genetically diverse populations. *PLoS genetics* **13,** e1006796 (2017).

5. Deredec, A., Burt, A. & Godfray, H. C. J. The population genetics of using homing endonuclease genes in vector and pest management. *Genetics* **179,** 2013–2026 (2008).

6. Harris, K. D., Hadari, G. & Greenbaum, G. modelRxiv: A platform for the distribution, computation and interactive display of models. *bioRxiv.* eprint: `https://www.biorxiv.org/content/early/2022/02/19/2022.02.16.480599.full.pdf` (2022).
